## Supplementary Information for "Viral fusion proteins of class II and III recognize and reorganize complex biological membranes"

**Table of Contents:**

Tables S1 to S7
Figures S1 to S10
Supplementary Results
Supplementary References

**Table S1:** Complex membrane mimicking the outer leaflet of the plasma membrane (*1*). In the second table, polyunsaturated tails were replaced with mono-unsaturated tails for PE and PS headgroup lipids. Lipid tail structures and their concentrations are listed in the table.

| Lipid | Structure | Conc. |
|---|---|---|
| POPC | 16:0-18:1 | 5% |
| PLPC | 16:0-18:2 | 5% |
| PAPC | 16:0-20:4 | 15% |
| DPSM | 18:1-18:1 | 10% |
| PLSM | 18:1-18:1 | 7% |
| PNSM | 18:1-18:1 | 8% |
| PAPE | 16:0-20:4 | 2% |
| PAPS | 16:0-20:4 | 4% |
| GM3 | 18:1-18:0 | 4% |
| CHOL | cholesterol | 40% |

| Lipid | Structure | Conc. |
|---|---|---|
| POPC | 16:0-18:1 | 5% |
| PLPC | 16:0-18:2 | 5% |
| PAPC | 16:0-20:4 | 15% |
| DPSM | 18:1-18:1 | 10% |
| PLSM | 18:1-18:1 | 7% |
| PNSM | 18:1-18:1 | 8% |
| POPE | 16:0-18:1 | 2% |
| POPS | 16:0-18:1 | 4% |
| GM3 | 18:1-18:0 | 4% |
| CHOL | cholesterol | 40% |

**Table S2:** Binary and ternary lipid membranes along with their structure and concentration. Lipid tail structures are shown for PC lipids in case of binary membranes, and for SM species and low abundant (5%) lipids in case of ternary membranes.

| Membrane | Structure | Conc. |
|---|---|---|
| DPPC/CHOL | PC 16:0-16:0 | 60%:40% |
| POPC/CHOL | PC 16:0-18:1 | 60%:40% |
| DOPC/CHOL | PC 18:1-18:1 | 60%:40% |
| PLPC/CHOL | PC 16:0-18:2 | 60%:40% |
| PAPC/CHOL | PC 16:0-20:4 | 60%:40% |
| PLPC/DPSM/CHOL | SM 18:1-16:0 | 30%:30%:40% |
| PLPC/PLSM/CHOL | SM 18:1-24:0 | 30%:30%:40% |
| PLPC/PNSM/CHOL | SM 18:1-24:1 | 30%:30%:40% |
| PLPC/PAPC/CHOL | PC 16:0-20:4 | 55%:5%:40% |
| PLPC/PAPE/CHOL | PE 16:0-20:4 | 55%:5%:40% |
| PLPC/PAPS/CHOL | PS 16:0-20:4 | 55%:5%:40% |
| PLPC/PAPI/CHOL | PI 16:0-20:4 | 55%:5%:40% |
| PLPC/GM3/CHOL | GM3 18:1-18:0 | 55%:5%:40% |

**Table S3:** Coarse-grained simulations to estimate the binding affinities of wild type and mutant (D961K) RVFV Gc to lipid membranes with and without PS lipids.

| Membrane | Structure | Conc. |
|---|---|---|
| PLPC/CHOL | PC 16:0-18:2 | 60%:40% |
| PLPC/PLPS/CHOL | PC/PS 16:0-18:2 | 48%:12%:40% |

**Table S4:** All-atom simulations of RVFV Gc and PrV gB. Final snapshots from coarse-grained simulations were backmapped to atomistic resolution for the systems listed in the table. Each system was simulated for 2 μs with 2 repeats.

| Membrane | Structure | Conc. |
|---|---|---|
| POPC/CHOL | PC 16:0-18:1 | 60%:40% |
| PAPC/CHOL | PC 16:0-20:4 | 60%:40% |
| PLPC/PAPE/CHOL | PC 16:0-18:2, PE 16:0-20:4 | 55%:5%:40% |
| PLPC/PAPS/CHOL | PC 16:0-18:2, PS 16:0-20:4 | 55%:5%:40% |
| PLPC/GM3/CHOL | PC 16:0-18:2, GM3 18:1-18:0 | 55%:5%:40% |

**Table S5:** Class I viral fusion proteins. The table lists the viral family, virus name, fusion protein, fusion peptide sequence, and uniportKB accession number. The full name of virus are given in the below description.

| Family | Virus | Fusion Protein | Fusion Peptide | UniProtKB Accession Number |
|---|---|---|---|---|
| Orthomyxoviridae | IAV | HA2 | GLFGAIAGFIENGWEGMIDG | P03442 |
| | IBV | HA2 | GFFGAIAGFLEGGWEGMIAG | P10757 |
| Retroviridae | HIV-1 | gp41 | AVGIGALFLGFLGAAGSTMGA | P03375 |
| | FIV | gp36 | VMLALATVLSIAGAGTGATAI | P16090 |
| | HTLV-1 | gp21 | AVPVAVWLVSALAMGAGVAGG | P23064 |
| | HTLV-2 | gp21 | AVPIAVWLVSALAAGTGIAGG | P03383 |
| | SIV | gp32 | GVFVLGFLGFLATAGSAMGAA | P05884 |
| | RSV | gp37 | GPTARIFASILAPGVAAAQAL | P03396 |
| | BLV | gp30 | VAALTLGLALSVGLTGINVAV | P51519 |
| Filoviridae | EBOV | gp2 | GAAIGLAWIPYFGPAAEGIYIEGLM | Q05320 |
| | MARV | gp2 | DLAAGLSWIPFFGPGIEGLYTAVLI | P35253 |
| Paramyxoviridae | SeV | F1 | FFGAVIGTIALGVATSAQITAGIAL | P04855 |
| | SV41 | F1 | FAGVVVGLAALGVATAAQVTAAVAV | P25181 |
| | MeV | F1 | FAGVVLAGAALGVATAAQITAGIAL | P69354 |
| | HeV | F | LAGVVMAGIAIGIATAAQITAGVAL | O89342 |
| | NDV | F | FIGAIIGSVALGVATAAQITAASAL | P26628 |
| | PIV5 | F | FAGVVIGLAALGVATAAQVTAAVAL | P04849 |
| Coronaviridae | HCoV-229E | S2 | SAIEDILFSKLVTSGLGTVDADYKKCTKGLSIADLACAQYYNG | P15423 |
| | HCoV-NL63 | S | SALEDLLFSKVVTSGLGTVDVDYKSCTKGLSIADLACAQYYNG | Q6Q1S2 |
| | FCoV | S | SAIEDLLFDKVVTSGLGTVDEDYKRCTGGYDIADLVCAQYYNG | P10033 |
| | SARS-CoV-2 | S2 | SFIEDLLFNKVTLADAGFIKQYGDCLGDIAARDLICAQKFNG | P0DTC2 |
| | SARS-CoV | S2 | SFIEDLLFNKVTLADAGFMKQYGECLGDINARDLICAQKFNG | P59594 |
| | MERS-CoV | S2 | SAIEDLLFDKVTIADPGYMQGYDDCMQQGPASARDLICAQYVAG | A0A023NGM8 |

Table S5 lists the class I viral fusion proteins and fusion peptide sequence used for conservation analysis as3 shown as matrix plot in Fig. 3. The representatives from Orthomyxoviridae family: IAV (Influenza A virus); IBV (Influenza B virus). Retroviridae family: HIV-1 (Human immunodeficiency virus type 1); FIV (Feline immunodeficiency virus); HTLV-1 (Human T-cell leukemia virus 1); HTLV-2 (Human T-cell leukemia virus 2); SIV (Simian immunodeficiency virus); RSV (Rous sarcoma virus); BLV (Bovine leukemia virus). Paramyxoviridae family: SeV (Sendai virus); SV41 (Simian virus 41); MeV (Measles virus); HeV (Hendra virus); NDV (Newcastle disease virus); PIV5 (Parainfluenza virus 5). Coronaviridae family: HCoV-229E (Human coronavirus 229E); HCoV-NL63 (Human coronavirus NL63); FCoV (Feline coronavirus); SARS-CoV-2 (Severe acute respiratory syndrome coronavirus 2 ); SARS-CoV (Severe acute respiratory syndrome coronavirus); MERS-CoV (Middle East respiratory syndrome-related coronavirus).

**Table S6:** Class II viral fusion proteins. The table lists the viral family, virus name, fusion protein, fusion loop sequence, and uniportKB accession number. The fusion loop hydrophobic aromatic residues inserting the membrane are underlined. The full name of virus are given in the below description.

| Family | Virus | Fusion Protein | Fusion Loop | UniProtKB Accession Number |
|---|---|---|---|---|
| Hantaviridae | HTNV | Gc | HCYGAC / YETS$\underline{W}$GCNPSDCPGVGTG / CN$\underline{F}$ATT | P08668 |
| | PUUV | Gc | HCYGSC / YETG$\underline{W}$GCNPPDCPGVGTG / CA$\underline{F}$ATT | P41266 |
| | DOBV | Gc | HCYGAC / YENN$\underline{W}$GCNPADCPGIGTG / CN$\underline{F}$ATT | Q806Y7 |
| | ANDV | Gc | HCYGAC / YETG$\underline{W}$GCNPGDCPGVGTG / CG$\underline{F}$ATT | Q9E006 |
| Togaviridae | CHIKV | E1 | VYP$\underline{F}$M$\underline{W}$GGAYC$\underline{F}$CDAENT | Q1H8W5 |
| | ONNV | E1 | VYP$\underline{F}$M$\underline{W}$GGAYC$\underline{F}$CDAENT | O90369 |
| | RRV | E1 | VYP$\underline{F}$M$\underline{W}$GGAYC$\underline{F}$CDSENT | P08491 |
| | SAGV | E1 | VYP$\underline{F}$M$\underline{W}$GGAYC$\underline{F}$CDTENT | Q9JGK8 |
| | SFV | E1 | VYP$\underline{F}$M$\underline{W}$GGAYC$\underline{F}$CDSENT | P03315 |
| | MAYV | E1 | VYP$\underline{F}$M$\underline{W}$GGAYC$\underline{F}$CDSENT | Q8QZ72 |
| | BFV | E1 | VYP$\underline{F}$M$\underline{W}$GGAYC$\underline{F}$CDTENS | P89946 |
| | AURV | E1 | VYP$\underline{F}$L$\underline{W}$GGAQC$\underline{F}$CDSENS | Q86925 |
| | SINV | E1 | VYP$\underline{F}$M$\underline{W}$GGAQC$\underline{F}$CDSENS | P03316 |
| | WEEV | E1 | VYP$\underline{F}$M$\underline{W}$GGAQC$\underline{F}$CDSENT | P13897 |
| | EEEV | E1 | VYP$\underline{F}$M$\underline{W}$GGAYC$\underline{F}$CDTENT | Q306W7 |
| Phenuiviridae | RVFV | Gc | CHLVGEC / CGG$\underline{W}$GCGC$\underline{F}$NVNP | P03518 |
| | PTV | Gc | CHLVGDC / CGAIGCGC$\underline{F}$NINP | P03517 |
| | CAV | Gc | CHLMSDC / CGAAGCGC$\underline{F}$NINP | F2W3Q1 |
| | TOSV | Gc | CHLMGEC / SGGIGYGC$\underline{F}$NVNP | A7LC97 |
| | MASV | Gc | CHLMGEC / SGGIGYGC$\underline{F}$NVNP | B6D3V7 |
| | SFSV | Gc | CRGVAEC / CGGVGCAC$\underline{F}$NVYA | Q88304 |
| | HTRV | Gc | CR$\underline{W}$AGDC / CGGAACGC$\underline{F}$NAAP | J3WAX0 |
| | SFTSV | Gc | CR$\underline{W}$AGDC / CGGAACGC$\underline{F}$NAAP | A0A0B5A886 |
| | UUKS | Gc | CHLMGAC / CGGALCQC$\underline{F}$NMRP | P09613 |
| Flaviviridae | DENV1 | E | RG$\underline{W}$GNGCGL$\underline{F}$G | P27912 |
| | DENV2 | E | RG$\underline{W}$GNGCGL$\underline{F}$G | P29990 |
| | DENV3 | E | RG$\underline{W}$GNGCGL$\underline{F}$G | P27915 |
| | DENV4 | E | RG$\underline{W}$GNGCGL$\underline{F}$G | Q2YHF2 |
| | TBEV | E | RG$\underline{W}$GNHCGL$\underline{F}$G | P14336 |
| | POWV | E | RG$\underline{W}$GNHCGF$\underline{F}$G | Q04538 |
| | WNV | E | RG$\underline{W}$GNGCGL$\underline{F}$G | Q9Q6P4 |
| | YFV | E | RG$\underline{W}$GNGCGL$\underline{F}$G | Q6DV88 |
| | ZIKV | E | RG$\underline{W}$GNGCGL$\underline{F}$G | A0A024B7W1 |

Table S6 lists the class II viral fusion proteins and fusion loop sequence used for conservation analysis as shown as matrix plot in Fig. 3. The residues from bc, cd and ij fusion loops are given for fusion proteins from Hantaviridae family; bc and cd fusion loop residues are given for fusion proteins from Phenuiviridae family and cd loop residues for Togaviridae and Flaviviridae families. The representatives from Hantaviridae family: HTNV (Hantaan virus); PUUV (Puumala virus); DOBV (Dobrava virus); ANDV (Andes virus). Togaviridae family: CHIKV (Chikungunya virus); ONNV (O'nyong-nyong virus); RRV (Ross river virus); SAGV (Sagiyama virus); SFV (Semliki forest virus); MAYV (Mayaro virus); BFV (Barmah forest virus); AURV (Aura virus); SINV (Sindbis virus); WEEV (Western equine encephalitis virus); EEEV (Eastern equine encephalitis virus). Phenuiviridae family: RVFV (Rift valley fever virus); PTV (Punta toro virus); CAV (Chandiru virus); TOSV (Toscana virus); MASV (Massilia virus); SFSV (Sandfly fever sicilian virus); HTRV (Heartland virus; SFTSV (Severe fever with thrombocytopenia syndrome virus)); UUKS (Uukuniemi virus). Flaviviridae family: DENV1 (Dengue virus type 1); DENV2 (Dengue virus type 2); DENV3 (Dengue virus type 3); DENV4 (Dengue virus type 4); TBEV (Tick-borne encephalitis virus); POWV (Powassan virus); WNV (West Nile virus); YFV(Yellow fever virus); ZIKV (Zika virus).

**Table S7:** Class III viral fusion proteins. The table lists the viral family, virus name, fusion protein, fusion loop sequence, and uniportKB accession number. The fusion loop hydrophobic aromatic residues inserting the membrane are underlined. The full name of virus are given in the below description.

| Family | Virus | Fusion Protein | Fusion Loop | UniProtKB Accession Number |
|---|---|---|---|---|
| Rhabdoviridae | VSIV | G | FRWYGP / CGYATV | P0C2X0 |
| | BEFV | G | ETWYFS / CFWNTE | P32595 |
| | VHSV | G | TSFFGG / IWMKNN | Q6QDI4 |
| | RABV | G | TNFVGY / HWVRTV | J7GA57 |
| Baculoviridae | AcMNPV | gp64 | YNGGSLDPNT / HFAH | P17501 |
| | BmNPV | gp64 | YNGGSLDPNT / HFAY | Q0EAF8 |
| | ApNPV | gp64 | YNGGPLDPNT / HFAH | Q1HH66 |
| | DisaGV | gp64 | YNGGSLDANT / HFAH | A0A0R7EYU3 |
| Orthomyxoviridae | THOV | gp | YNGGLVDSNT / KFAY | P28977 |
| | DHOV | gp | YNGGSLDKNT / HFAY | P27427 |
| | SINUV | gp | YQGGPLDPNT / HFAH | A0A1L5YKF7 |
| | BRBV | gp | YNGGSLDGNT / HFAH | A0A140H4W8 |
| Herpesviridae | HSV1 | gB | VWFGHRY / RVEAFHR | P06437 |
| | HSV2 | gB | VWFGHRY / RVEAFHR | P06763 |
| | PrV | gB | VWSGSTY / GAAGFYH | G3G8X1 |
| | VZV | gB | AWAGSSY / GTPGTYR | P09257 |
| | HCMV | gB | SYAYIYT / GSTWLYR | P06473 |
| | EBV | gB | IYNGWYA / GWLIWTYR | P03188 |

Table S7 lists the class III viral fusion proteins and fusion loop sequence used for conservation analysis as shown as matrix plot in Fig. 3. The residues from fusion loops I and II are given for fusion proteins. The representatives from Rhabdoviridae family: VSIV (Vesicular stomatitis indiana virus); BEFV (Bovine ephemeral fever virus); VHSV (Viral hemorrhagic septicemia virus); RABV (Lyssavirus rabies). Baculoviridae family: AcMNPV (Autographa californica nuclear polyhedrosis virus); BmNPV (Bombyx mori nuclear polyhedrosis virus); ApNPV (Antheraea pernyi nuclear polyhedrosis virus); DisaGV (Diatraea saccharalis granulovirus). Orthomyxoviridae family: THOV (Thogoto virus); DHOV (Dhori virus); SINUV (Sinu virus); BRBV (Bourbon virus). Herpesviridae family: HSV1 (Human herpesvirus 1); HSV2 (Human herpesvirus 2); PrV (Pseudorabies virus); VZV (Varicella-zoster virus); HCMV (Human cytomegalovirus); EBV (Epstein-Barr virus).

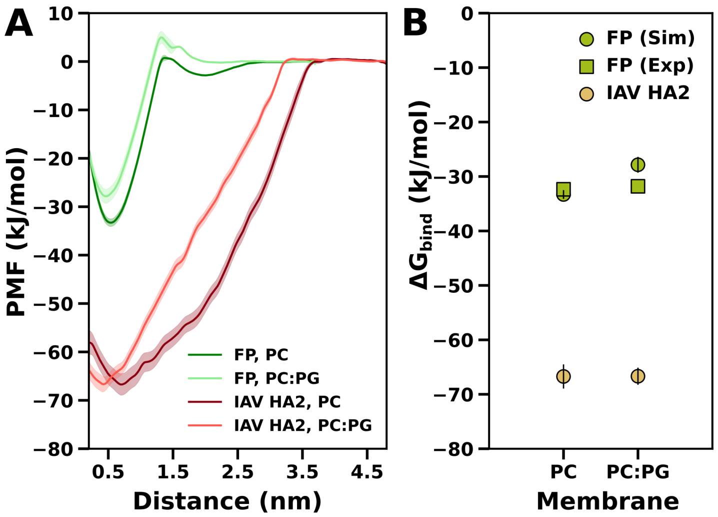


**Figure S1: Binding affinity calculations of the full-length IAV HA2 trimer and of the fusion peptide (FP) monomer to lipid membranes**. (**A**) Potentials of mean force (PMFs) for IAV HA2 trimer (red shades) and FP monomer (green shades) binding to membranes of pure POPC (PC) or POPC:POPG (PC 80%:PG 20%, see legend). (**B**) Binding free energies $\Delta G_{\text{bind}}$ as taken from PMFs in panel (A) for IAV HA2 (brown circles) and FP (green circles) in PC and PC:PG membranes. For comparison, experimental $\Delta G_{\text{bind}}$ values for the FP are shown as green squares (*3*). Experiments revealed, for binding of a single IAV HA2 fusion peptide to POPC:POPG (80%:20%), $\Delta G_{\text{bind}}$ of −31.8 kJ/mol at pH 5.0 and of −33.7 kJ/mol at pH 7.4 (*3*). For a POPC membrane, the experimental $\Delta G_{\text{bind}}$ was −32.35 kJ/mol at pH 5.0 and −30.30 kJ/mol at pH 7.4 (*3*). The IAV HA2 fusion peptide structure used in our simulations corresponds to the NMR structure solved at pH 5.0 (PDB ID: 1IBN) (*4*), hence we compare the $\Delta G_{\text{bind}}$ measured at pH 5.0 with our simulations. Excellent agreement is found between simulation and experiments (compare green circles with green squares). Notably, the binding affinity of the trimer (brown circles) is not three times the affinity of the FP monomer (green circles), possibly because binding of the entire fusion protein involves a larger perturbation of the membrane as compared to binding of only three FPs, thereby rendering the binding less favorable. Shaded areas in (A) and vertical black bars in (B) denote 1 SE

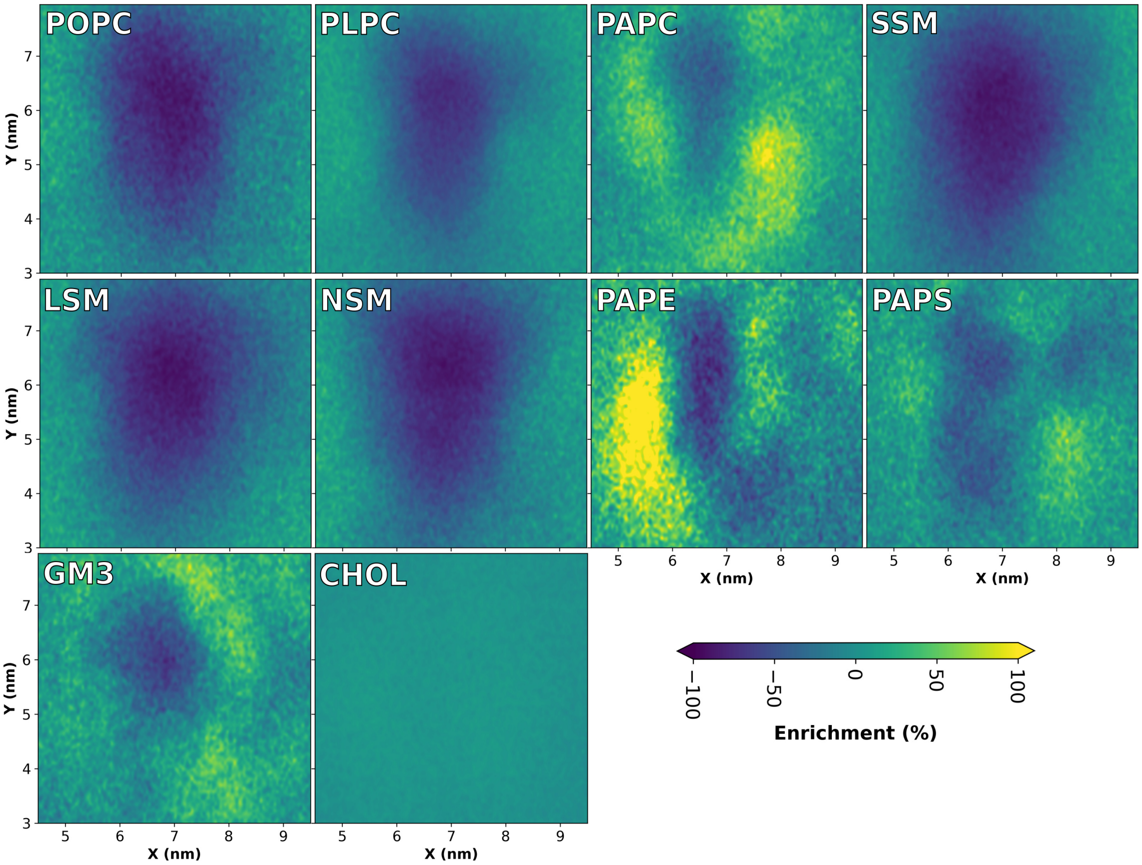


**Figure S2: Lateral lipid densities in coarse-grained simulation of IAV HA2 bound to the plasma membrane model**. Two-dimensional density map of each lipid bound IAV HA2, plotted as the enrichment relative to the density in the bulk membranes. Although polyunsaturated lipids (PAPC, PAPE, yellow areas) and GM3 (light green areas) are enriched near the fusion peptides, no well-defined binding sites are found.

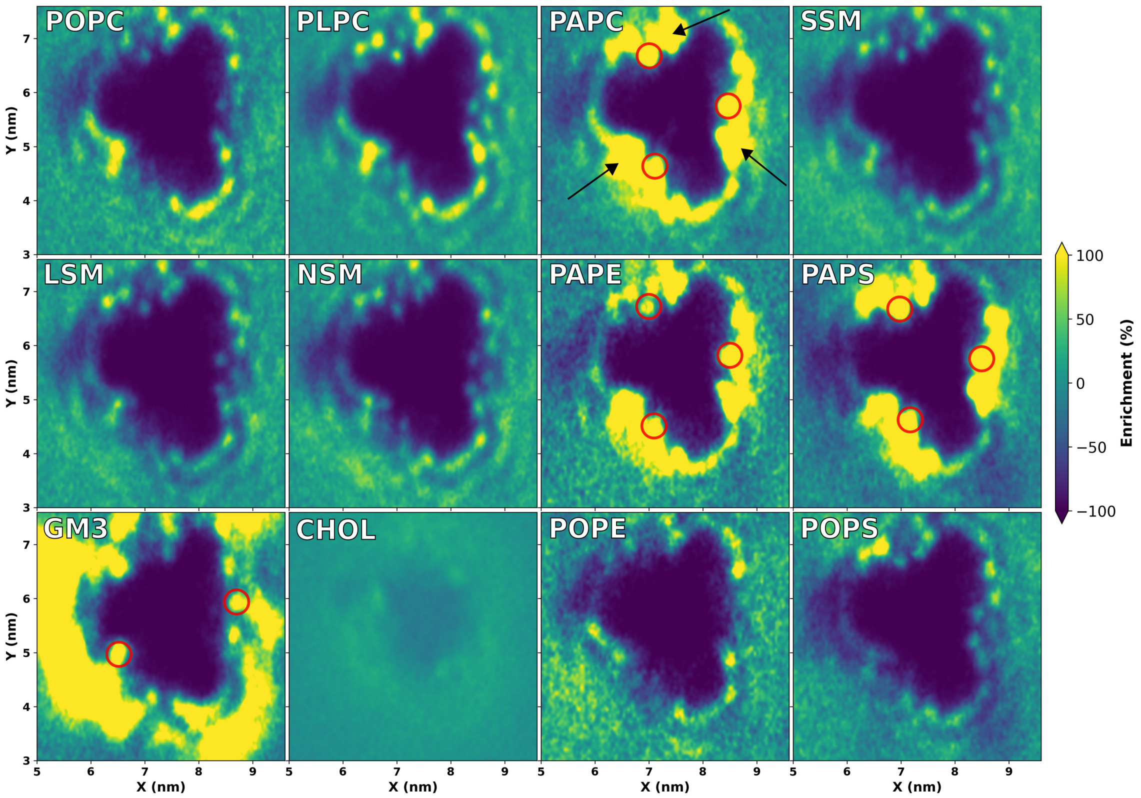


**Figure S3: Lateral lipid densities in coarse-grained simulation of PrV gB bound to the model of the outer leaflet of the plasma membrane**. Two-dimensional density map of each lipid bound PrV gB, plotted as the enrichment relative to the bulk membranes. Lipid binding sites at the monomer–monomer interface are marked in red circles. Polyunsaturated lipids (PAPC, PAPE, PAPS) as well as GM3 are greatly enriched at the protein.

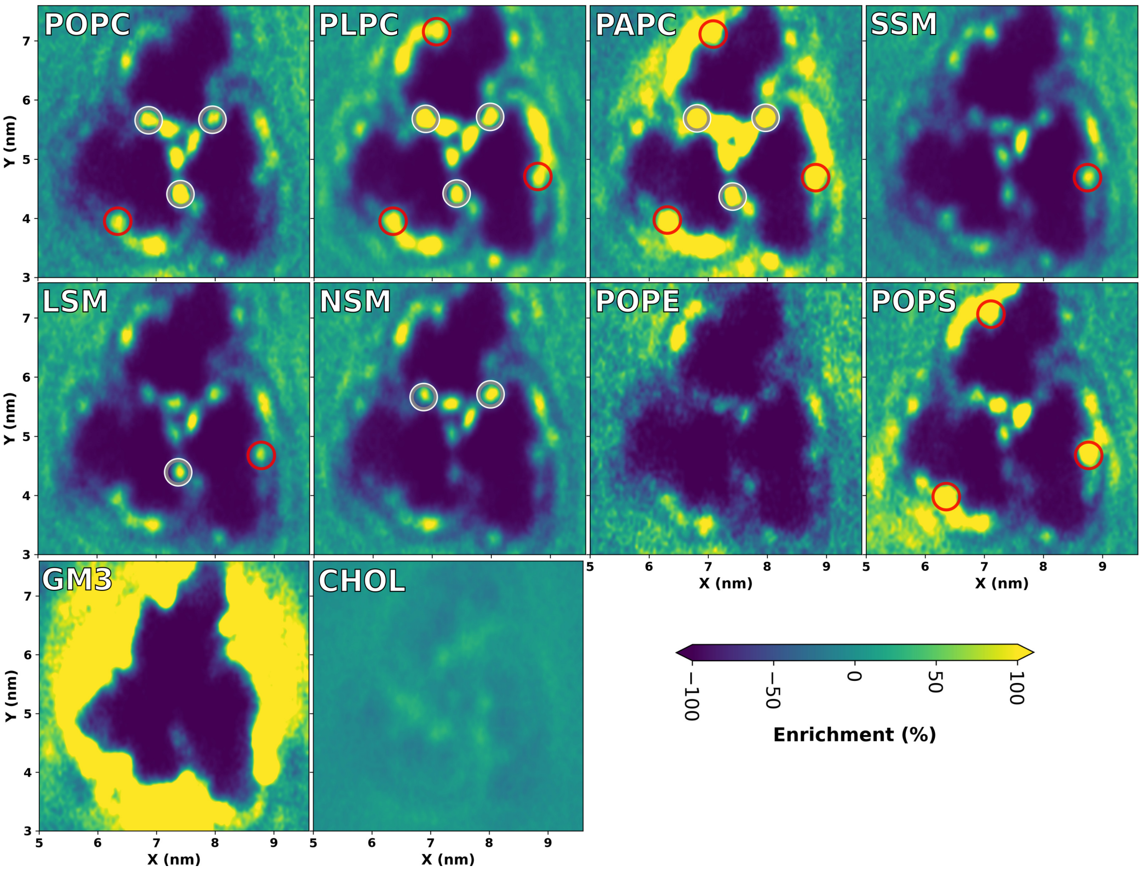


**Figure S4: Lateral lipid densities in coarse-grained simulations of RVFV Gc bound to a plasma membrane model containing monounsaturated POPE, POPS lipids**. Two-dimensional density map of each lipid bound RVFV Gc, plotted as the enrichment relative to the bulk membranes. Lipid binding sites on each monomer and at the monomer–monomer interface are marked with red and grey circles respectively. Polyunsaturated PAPC as well as GM3 are greatly enriched at the protein.

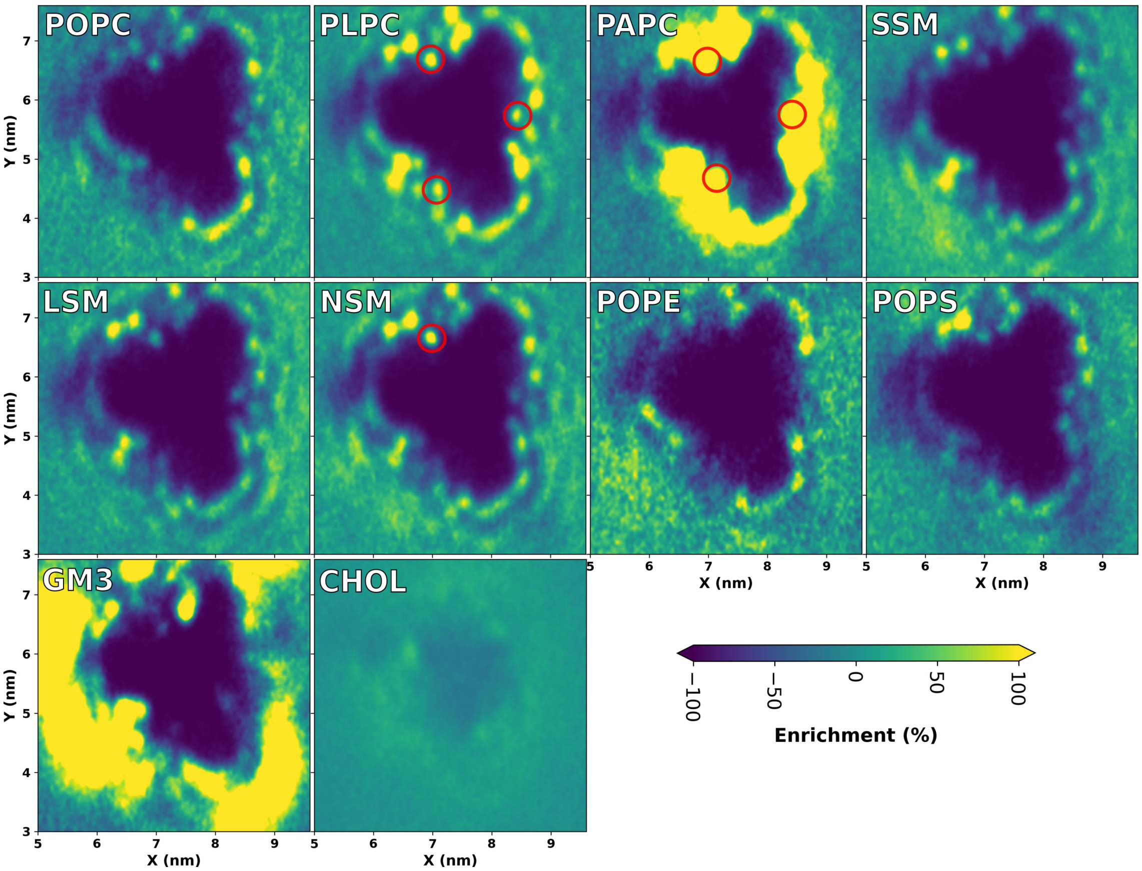


**Figure S5: Lateral lipid densities in simulations of PrV gB bound to a plasma membrane model containing monounsaturated POPE, POPS lipids**. Two-dimensional lipid density maps of lipids around PrV gB, plotted as the enrichment relative to the density in the bulk membranes. Lipid binding sites at the monomer–monomer interface are marked in red circles. Polyunsaturated PAPC as well as GM3 are greatly enriched at the protein.

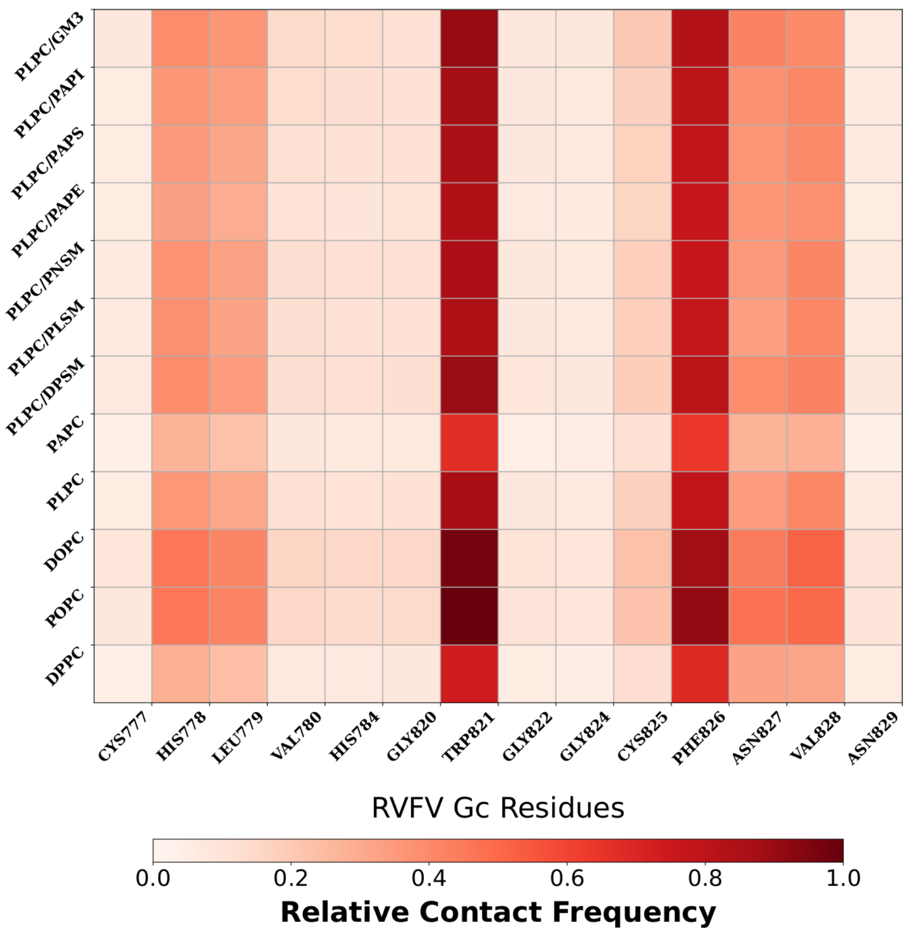


**Figure S6: Interaction of cholesterol with RVFV Gc residues in binary and tertiary lipid membranes**. All systems contain 40% cholesterol plus two additional phospholipids (upper seven rows) or plus one additional phospholipd (lower five rows). Contact frequencies are normalized to 100% by dividing the contacts in each system by cholesterol concentration (40%). Cholesterol–protein distances within 0.65 nm were defined as contact. Here, cholesterol interacted primarily with residue Trp821 and Phe826.

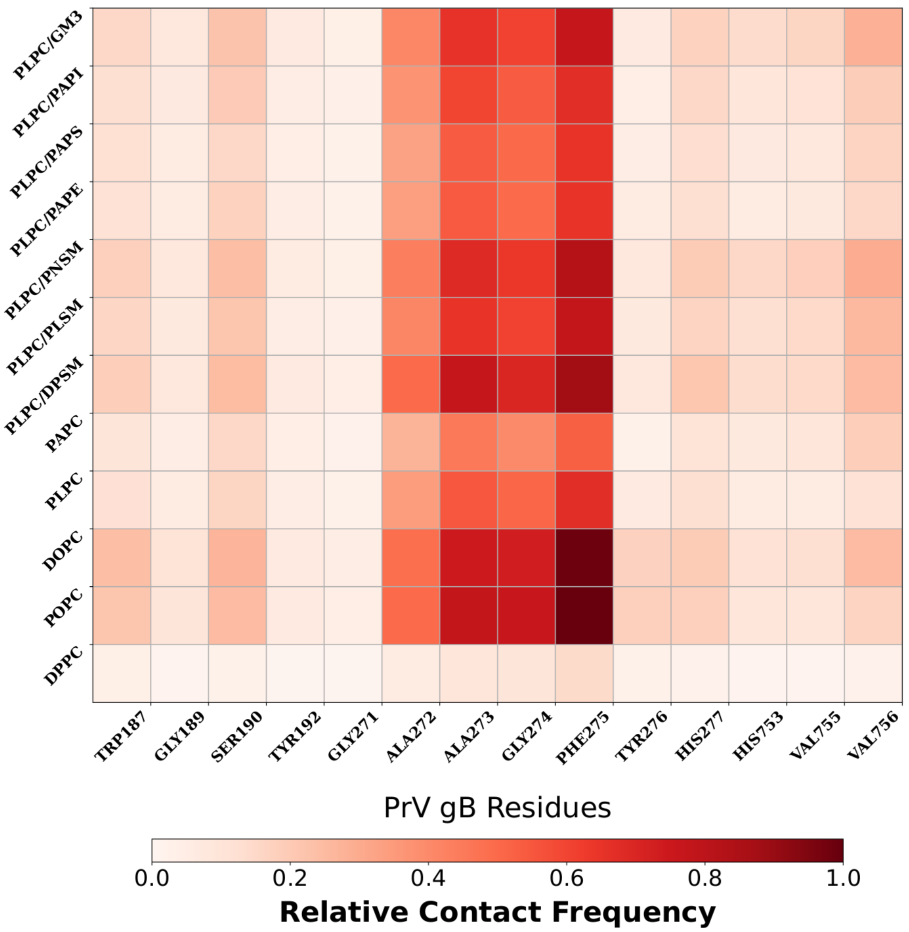


**Figure S7: Interaction of cholesterol with PrV gB residues in binary and tertiary lipid membranes**. All systems contain 40% cholesterol plus two additional phospholipids (upper seven rows) or plus one additional phospholipd (lower five rows). Contact frequencies are normalized to 100% by dividing the contacts in each system by cholesterol concentration (40%). Cholesterol–protein distances within 0.65 nm were defined as contact. Here, cholesterol interacted primarily with residue Phe275.

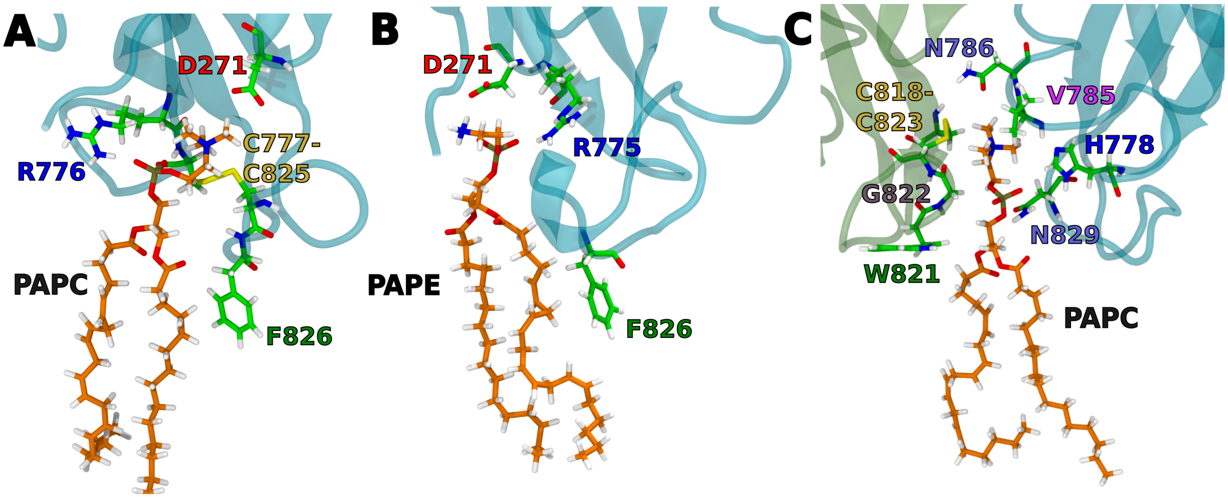


**Figure S8: Snapshots from all-atom simulations depict the lipid binding site on the monomer (A,B) and at the monomer–monomer interface (C) in RVFV Gc**. RVFV Gc do not distinguish between polyunsaturated PC and PE headgroups as indicated by similar $\Delta G_{\text{bind}}$ (Fig. 2D). The similar binding for PC and PE lipids in our simulations is in agreement with bio-layer interferometry (BLI) experiments which showed that PE lipids restore RVFV Gc binding when added to liposomes composed of SM and cholesterol (*5*). Atomistic simulations further corroborate the experimental findings, as we observe PE lipids to occupy the same binding site as PC lipids (A,B).

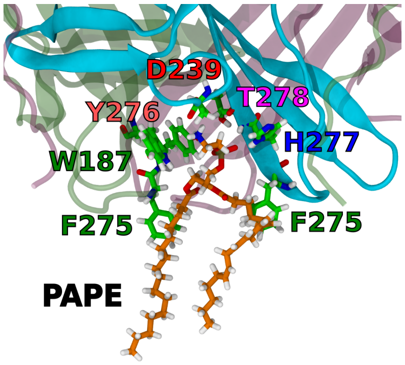


**Figure S9: Snapshot from all-atom simulation depict the polyunsaturated PE lipid binding at monomer–monomer interface in PrV gB**. PrV gB do not distinguish between polyunsaturated PC and PE headgroups as indicated by similar $\Delta G_{\text{bind}}$, (Fig. 2D) The observed similar binding for PC and PE lipids is in agreement with liposome flotation assay on gB (*6*).

### Supplementary Results

##### Fully saturated sphingomyelin (SM) reduces binding, whereas unsaturated SM does not interfere with binding

Along with PC lipids, sphingomyelin (SM) is a highly abundant phospholipid in the outer leaflet of the PM. Binding experiments have shown weak binding or complete loss of binding by RVFV Gc to liposomes containing palmitoyl sphingomyelin (PSM) or PSM/cholesterol, respectively (*5*). Recent lipidomics data has shown that the outer leaflet of the PM contains a large fraction of SM with long or unsaturated chain fatty acids such as 18:1-24:0 or 18:1-24:1 (*1*).

To investigate how SM affects binding free energies, we set up systems with varying SM tail length and unsaturation (Fig. 2D). Addition of palmitoyl sphingomyelin (PSM, 18:1-16:0) to the reference lipid mixture of PLPC/cholesterol reduces the binding affinity (increases $\Delta G_{\mathrm{bind}}$) in line with experimental binding data (*5*). However, we found that weak binding is recovered upon increasing the tail length or unsaturation from C16:0 to C24:0 (Lignoceroyl SM, LSM) or C24:1 (Nervonoyl SM, NSM). Similar effects are found for IAV HA2 binding. For PrV gB, in contrast, the SM tail length and unsaturation have only a minor effect on binding, in line with experiments showing that PrV gB binding strictly requires 40% cholesterol but does not depend on SM or other lipid content (*6*). We explain the observed difference in binding upon addition of SM species by the varied membrane-stiffening properties of SM species, which follow order of PSM (16:0) $>$ LSM (24:0) $>$ NSM (24:1) (*7, 8*).

##### Anionic PS lipids rescue membrane binding of mutant D961K Gc

Previous experiments investigated the effect of a mutation in the lipid binding pocket of RVFV Gc on membrane binding (*5*). Specifically, mutating the choline-interacting aspartic acid D961 to lysine (Fig. 5D) resulted in loss of binding to PC/cholesterol membranes due to electrostatic repulsion between the cationic lysine and choline moieties. However, binding was rescued upon the addition of PS lipids, which was attributed to favorable interactions between anionic PS and cationic lysine residue (*5*). These findings offer insight into the lipid specificity of class II fusion proteins.

Consistent with the experimental results, the wild-type Gc exhibits similar affinity for membranes with and without PS lipids, indicating that PS lipids are not significant for wild type binding (Fig. 2B-D, Fig. S10). Visual inspection of the simulations confirms that the similarity in affinity for PS lipids stems from their deep insertion into the binding site, where the carboxyl and phosphate groups form hydrogen bonds with the side chains of R775 and R776, respectively (Fig. 5E). In sharp contrast, the D961K mutant displays 20 kJ/mol stronger affinity ($\Delta G_{\mathrm{bind}}$ more negative) for PS-containing membranes compared to PS-free membranes, corresponding to a approximately $\sim$3000-fold increased binding probability (Fig. S10). The recovery of binding by the D961K mutant upon the addition of PS lipid results from the localization of PS lipid at the binding site. The lipid-sensitivity of residue 961 found in both simulation and experiments confirms the importance of the PC binding site.

Alongside PS, the endosomal membranes with which class II viral proteins interact are rich in other anionic lipids such as phosphatidylinositol (PI) and Bis(monoacylglycero) phosphate (BMP) (*9–14*). To investigate the potential role of phosphatidylinositol (PI) lipids in membrane binding, we substituted the phosphatidylcholine (PC) headgroup with PI. We observed that the wild type Gc binds strongly to the anionic headgroup of PI (Fig. 2 D), likely facilitated by additional electrostatic interactions involving cationic residues near the lipid binding pockets. Thus, the presence of PI lipids enhances binding, aligning with the virus's entry via the endocytic pathway (*9, 15*).

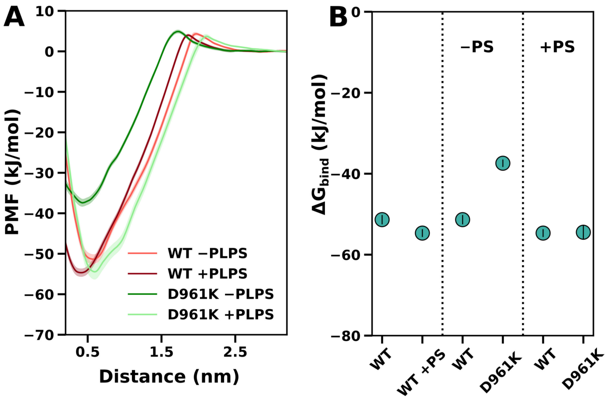


**Figure S10: Effect of the D961K mutation on RVFV Gc binding to membranes**. (**A**) Potentials of mean force (PMFs) for wild type (WT) and D961K mutant of RVFV Gc binding to PLPC/cholesterol/(+/− PLPS) membranes. (**B**) Binding free energies taken from the PMFs for WT and D961K RVFV Gc binding to PLPC/cholesterol/(+/− PLPS) membranes. Shaded areas in (A) and vertical black bars indicate 1 SE.
